# Neolactotetraosylceramide: A novel non-invasive urinary biomarker for bladder cancer

**DOI:** 10.1101/2023.08.08.552442

**Authors:** Inês B. Moreira, Charlotte Rossdam, Julia Beimdiek, Manuel M. Vicente, Jessica Schmitz, Astrid Oberbeck, Jan H. Bräsen, Hossein Tezval, Falk F. R. Buettner

## Abstract

There is an urgent need to identify noninvasive biomarkers for bladder cancer. Here, we applied glycan analytics by multiplex capillary gel electrophoresis coupled to laser-induced fluorescence detection (xCGE-LIF) to unravel the global glycosphingolipid (GSL)-glycan profile of primary tumor tissues and urine samples from bladder cancer patients. Thereby, we detected neolactotetraosylceramide (Galβ1-4GlcNAcβ1-3Galβ1-4Glc-Cer, nLc4) at significantly increased levels from tumorigenic regions of bladder tissues compared to non-malignant adjacent material (n = 30). Specific expression of nLc4 in cancer tissue was confirmed by immunofluorescence staining. GSL-glycan profiling by xCGE-LIF of urinary exosomes showed that nLc4 is increased in bladder cancer patients (n = 16) when compared to controls (n = 50), with an overall sensitivity of 57% and specificity of 90%. We set-up an ELISA targeting nLc4-containing urinary exosomes from bladder cancer patients (n = 9) and cancer-free individuals (n = 9) demonstrating an overall sensitivity and specificity of 89% and 78%, respectively.

**SIGNFICANCE:** This study shows that levels of nLc4 are significantly elevated in bladder cancer tissue and urinary exosomes of bladder cancer patients. Urinary detection of nLc4 by glycan analytics or ELISA outperforms standard diagnostic modalities, facilitating noninvasive bladder cancer diagnosis. Furthermore, nLc4 bears the potential of being a target for bladder cancer therapy.

## INTRODUCTION

Bladder cancer (BC) is the most common malignant tumor in the urinary system, being transitional cell carcinoma its main pathological feature. Worldwide, BC is the tenth general most common cancer and accounted for more than 573 000 new cases and 212 000 deaths in 2020 (1). Currently, the diagnosis of BC is based on cystoscopic examination and histological evaluation of the tissue, usually done in patients already with signs or symptoms of it. Being an invasive procedure that can cause complications like bleeding or urinary tract infections, cystoscopy is often accompanied by urine cytology, a test that screens for exfoliated tumor cells in the urine. Although representing a simple and non-invasive method, urine cytology lacks adequate sensitivity and overall efficacy to be used as the primary method of histological diagnosis, especially for low-grade BC (2–6). Moreover, while more than 30 urinary biomarkers have been recognized so far to diagnose BC, only a few have formal indication for clinical practice (7, 8). After diagnosis, treatment options, which differ according to cancer type and stage, conventionally include surgery, cystectomy and chemo-, radio-and immunotherapy (9, 10). However, these modalities encompass an increased economic burden, as well as a need for targeted delivery. Subsequently, there is an ongoing effort to identify new reliable biomarkers in urine and tissue, as accessible alternative tools for the early diagnosis and treatment of BC.

Overall, tumor cells display a set of oncogenic-related cellular modifications that confers them selective advantage. Among them, alterations in glycosylation pathways are a common feature of all cancer hallmark abilities, with most FDA-approved tumor markers being glycan antigens or glycoproteins (11). These carbohydrate structures, which are assembled in a stepwise manner by the coordinated action of different glycosyltransferase and glycosidase enzymes, can be attached to proteins or lipids. Glycosphingolipids (GSLs), glycoside compounds that consist of a hydrophobic ceramide backbone decorated with a hydrophilic carbohydrate residue, are the major glycolipids of mammals (12). These glycans, mainly classified within ganglio-, globo-, or (neo)lacto-series, are known to interact with key molecules at the cell membrane level, playing essential roles in mediating cell-cell interactions and modulating signal transduction pathways (12, 13). Aberrant expression of specific GSLs is strongly associated with tumor initiation and malignant transformation in many types of human cancer, altering cell growth, adhesion and motility (14). As different cells and tissues show differential expression of GSLs, BC specific alterations can be identified by multiplex gel-electrophoresis coupled to laser-induced fluorescence detection (xCGE-LIF), a fast and high-throughput-compatible glyco-analytical approach that enables the screening of potential glycan markers derived from complex biological samples (15).

In this study, we applied glycan analytics by xCGE-LIF for quantitative GSL-glycan profiling of primary BC tumors and non-malignant surrounding material as well as urine samples of BC patients and non-BC individuals. To our knowledge, this approach unraveled for the first time the global GSL profile of bladder cancer cohorts, which led to the identification of nLc4 as a new biomarker for BC.

## RESULTS

### Glycosphingolipid-glycan analysis of bladder cancer tissue uncovered tumor-specific signatures

We quantitatively profiled glycosphingolipid-derived glycans by xCGE-LIF from formalin fixed paraffin embedded (FFPE) tissue samples of the primary tumor (bladder cancer, BC) and non-malignant surrounding material (normal adjacent tissue, NAT) from 30 patients (cohort 1). Moreover, bladder tissues from 7 cancer-free (CF) individuals were included. Clinical data related to the applied samples are available in Supplementary Tables S1 and S2. A workflow schematically displaying sample processing and analysis is provided as Supplementary Fig. S1. Structures of all GSLs identified in this study are available in Supplementary Table S3. Overall, 13 different GSL species could be detected in the analyzed tissue samples by xCGE-LIF, of which 9 species (GM3, GD3, GD1b, GA1, Gb3, Gb4-like, nLc4, α6-sialyl nLc4 and galabiose) are exclusively found in cancer patients, but not in cancer-free bladder material (Fig. 1A, Supplementary Tables S4 and S5). From those cancer patient-specific glycans, GM3, Gb3, Gb4-like, nLc4 and α6-sialyl nLc4 are significantly increased in BC when compared to the correspondent NAT (Fig. 1A, Supplementary Fig. S2). Principal component analysis (PCA) based on relative GSL abundance reveals that the cluster comprising CF samples is well separated from the BC and - with one exception - from the NAT clusters. NAT samples also cluster well whereas BC samples are more broadly distributed and show overlap with the NAT samples (Supplementary Fig. S3A). In the following we focus on GM3, Gb3 and nLc4 which are the glycan precursors of the ganglio-, globo-and neolacto-series, respectively. In this context, we evaluated the correlation between precursors and their derivatives’ expression and found that Gb4-like (derivative of Gb3) and α6-sialyl nLc4 (derivative of nLc4) are positively correlated to their precursors in the cancer samples (Supplementary Fig. S3B). GM3 levels are increased in 25 out of 30 BC vs. NAT sample pairs (mean fold change: 3.1, *P* < 0.0001, Fig. 1B), Gb3 levels are increased in 24 out of 30 BC vs. NAT sample pairs (mean fold change: 2.4, *P* < 0.0001, Fig. 1C) and nLc4 is present in 11 BC samples but not in any of the NAT or CF samples (fold change could not be calculated as all control values were 0, *P* = 0.0010, Fig. 1D). We used receiver operating characteristic (ROC) curve analysis to assess the diagnostic efficacy of GM3, Gb3 and nLc4 (Fig. 1B-D) and all other detected GSLs (Supplementary Fig. S4). Among the three series precursor glycans, GM3 shows the highest area under the ROC curve value (AUC = 0.851, 95% confidence interval (CI) = 0.758-0.944, *P* < 0.001), followed by Gb3 and nLc4. For each marker, the cut-off point in terms of relative signal intensity was defined at the highest Youden’s index (sensitivity + specificity – 1) (Supplementary Table S6), reaching optimal sensitivity and specificity values of 73.3% and 86.5% for GM3, 76.7% and 70.3% for Gb3 and 36.7% and 100% for nLc4, respectively. Logistic regression was used to build a diagnostic model that could explore whether combinations of biomarkers could improve the diagnostic efficacy for patients with BC. The combination of GM3, Gb3 and nLc4 resulted in a superior AUC (0.885, 95% CI= 0.821-0.949, *P* < 0.001; Fig. 1E), which was higher than the AUC values for each marker alone. The combined diagnostic model reached 73.3% of sensitivity and 94.6% of specificity. There are no significant differences for the identified GSLs regarding the histological grade or pathological stage such as non-muscle invasive or muscle invasive bladder cancer (Supplementary Fig. S5).

**Figure 1.**
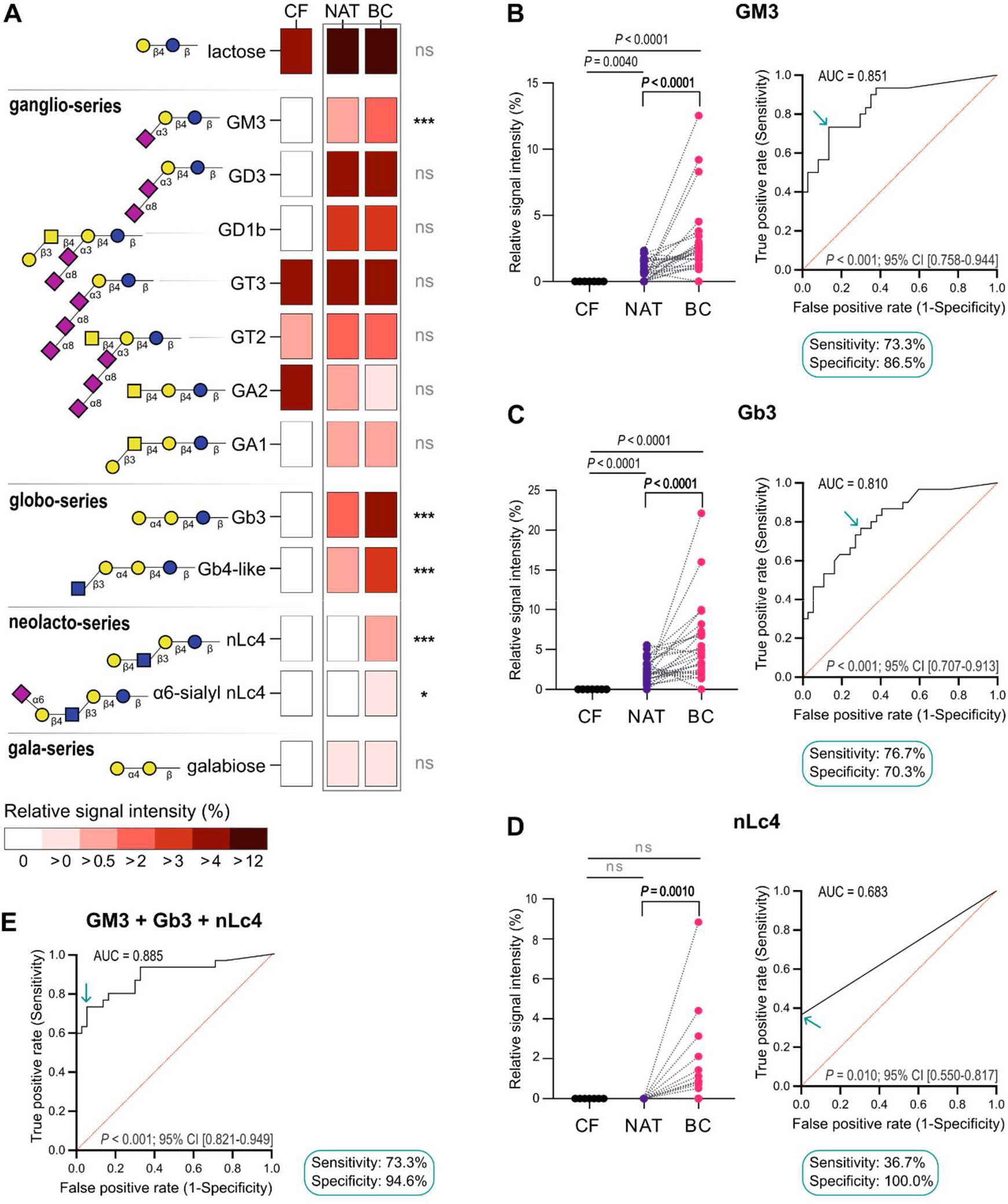
Glycosphingolipids detected in bladder cancer (BC, n = 30), normal adjacent tissue (NAT, n = 30) and cancer-free (CF, n = 7) tissue samples through xCGE-LIF. **A,** Heatmap showing relative signal intensity mean values of 13 GSLs identified in BC. *Q* values were calculated using Multiple Mann-Whitney test (false discovery rate of 5%). *, *Q* value <0.05; **, *Q* value <0.005; ***, *Q* value <0.001; ns, non-significant. **B – D,** Subset of the GSLs that are overexpressed in cancer and undetectable in CF samples. Levels of **B,** GM3, **C,** Gb3 and **D,** nLc4 in CF and paired analysis between NAT and BC (left). *P* values were calculated using two-tailed unpaired Mann-Whitney test (for comparisons with the CF group) or two-tailed Wilcoxon matched-pairs signed rank test (for NAT *vs.* BC comparison); Receiver operating characteristic (ROC) curve analysis in bladder cancer detection obtained by calculating the sensitivity and specificity of the test at every possible cut-off point and plotting the sensitivity against 1-specificity are shown on the right. **E,** ROC curve analysis obtained through logistic regression of GM3, Gb3 and nLc4 markers combined. Area under the curve (AUC), *P* values and 95% confidence intervals (CI) are shown for each ROC curve. The arrows indicate the cut-off values of relative signal intensity that best discriminate BC detection, for which the correspondent sensitivity and specificity values are shown. Blue circle: glucose, yellow circle: galactose, blue square: *N*-acetylglucosamine, yellow square: *N*-acetylgalactosamine, pink diamond: *N*-acetylneuraminic acid.

In order to further investigate GSL-glycosylation associated with BC, we additionally performed an *in-silico* analysis of genes that participate in the GSL biosynthesis pathway using a publicly available gene expression bladder cancer dataset (TCGA-BLCA, The Cancer Genome Atlas Urothelial Bladder Carcinoma; https://portal.gdc.cancer.gov/projects/TCGA-BLCA) (16). The expression of *A4GALT* (Gb3 synthase) and *ST3GAL5* (GM3 synthase) genes is downregulated in BC when compared to the NAT group which does not correlate to our findings by glycan analytics. Interestingly, expression of *B4GALT3* and *B4GALT4* which encode glycosyltransferases that are directly involved in the biosynthesis of nLc4, is upregulated. Moreover, *ST6GAL1*, which encodes the glycosyltransferase that generates α6-sialyl nLc4 through the addition of a sialic acid to nLc4, is downregulated in BC, which can contribute to the accumulation of the nLc4 precursor in BC tissues as well (Supplementary Fig. S6). Based on these findings, we focused on nLc4 as our major glycan target.

### nLc4 is specifically expressed on bladder cancer tissue

As neither xCGE-LIF nor gene expression analyses provide information on the spatial distribution of glycans at tissue level, we combined hematoxylin and eosin staining (H&E-staining) with multiplexed immunohistochemistry on identical tissue sections to analyze the expression of nLc4 and GATA3 in BC and CF bladder tissues (Fig. 2). H&E staining clearly shows morphological alterations between bladder cancer and control tissue (Fig. 2A). nLc4 expression is broadly distributed over the tumor in the BC tissue sample but not detected in the CF sample (Fig. 2B). Both samples were additionally stained against GATA3, which is a valuable marker of urothelial differentiation expressed in benign cells of the urothelium but also in derived urothelial carcinoma (17). In the control tissue, the normal mucosa shows strong nuclear GATA3 positivity (Fig. 2B, control). On the other hand, upon urothelial neoplasm, GATA3-expressing cells get diffusely distributed throughout the tumor tissue (18) as observed in our BC tissue samples (Fig. 2B). Notably, our analyses show that nLc4 and GATA3 staining colocalize within the tumor regions (Fig. 2B).

**Figure 2.**
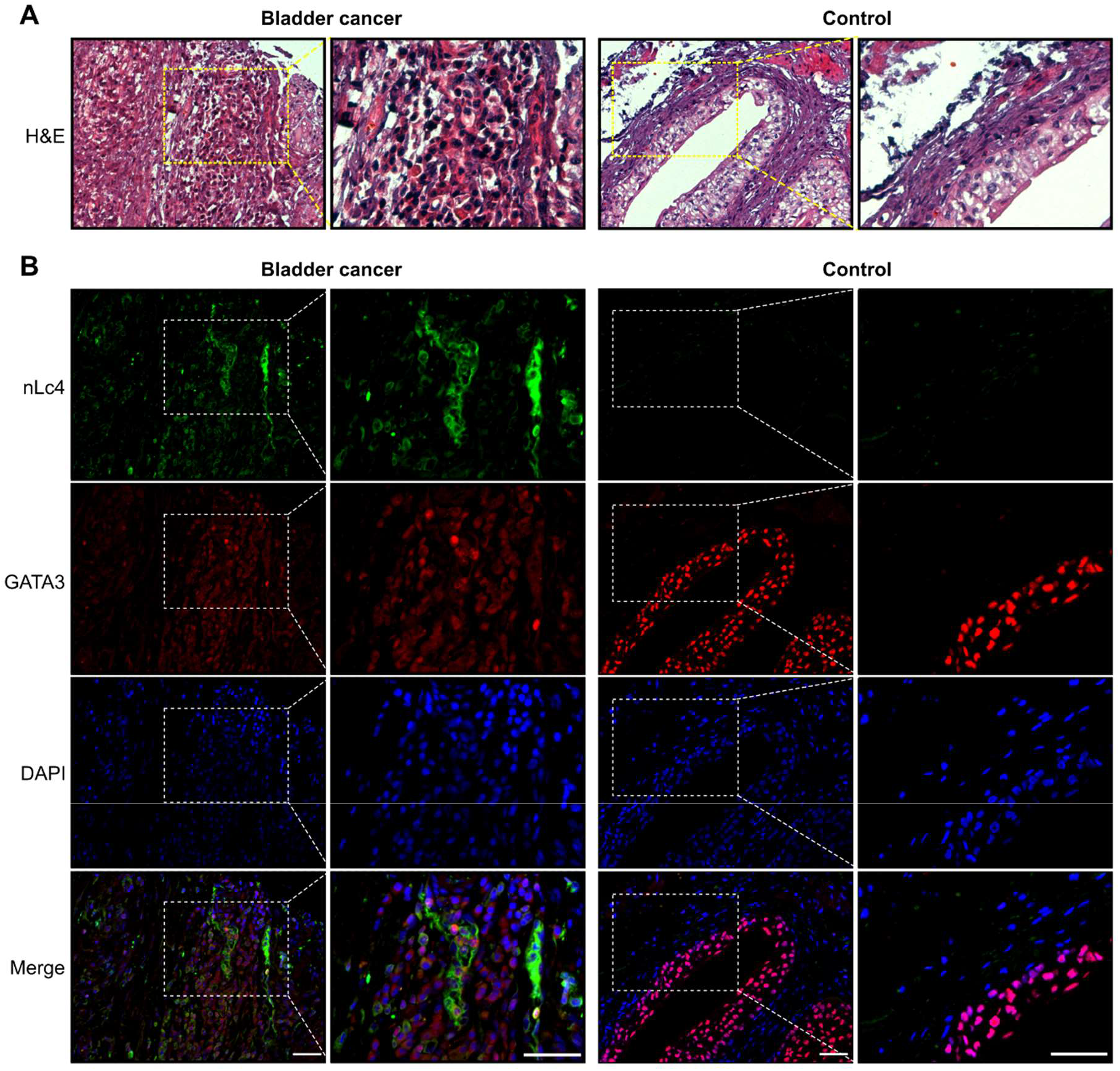
Multiplex immunofluorescence (mIF) and haematoxylin & eosin (H&E) staining on identical tissue sections of BC and control, respectively. Representative **A,** H&E and **B,** mIF staining images of BC and control tissues labeled with nLc4 (Clone 1B2) (green) and GATA3 (red). DAPI was used to visualize nucleus (blue). Scale bars correspond to 50 μM.

To ensure that the anti-nLc4 antibody (1B2) is specific for the glycosphingolipid nLc4 (Galβ1- 4GlcNAcβ1-3Galβ1-4Glc-Cer) and does not cross-react with *N*- or *O*-glycans terminating with similar structures, we generated a BC cell line completely lacking glycosphingolipids to be tested for detection by 1B2. Therefore, we introduced a frameshift mutation into the UDP-glucose ceramide glycosyltransferase (*UGCG*) gene encoding the glucosylceramide synthase (GCS) in the human BC cell line (CAL-29) by CRISPR/Cas9. xCGE-LIF analysis as well as antibody-and lectin-staining of the generated CAL-29 *UGCG*-knockout (KO) cell line revealed complete absence of glycosphingolipids while *N*-glycans terminating with Galβ1-4GlcNAc could still be detected by ECL lectin (data not shown). Analysis of CAL-29 wildtype (WT) and *UGCG*-KO cells by immunocytochemistry and flow cytometry revealed that the 1B2 antibody detects the wildtype but not the knockout cells strongly suggesting its specificity towards the GSL nLc4 (Supplementary Fig. S7). Taken together, our analyses confirm that nLc4 is specifically expressed on BC tissue but not on healthy urothelium.

### Urinary glycosphingolipid profiling suggests nLc4 as a non-invasive biomarker for bladder cancer

Motivated by the need for the identification of novel non-invasive biomarkers of BC, we set out to apply xCGE-LIF for urinary GSL-glycan profiling. Therefore, we analyzed GSLs by xCGE-LIF of extracellular vesicles (EVs) prepared from urine samples (Supplementary figure S1) of 16 patients with BC and 50 non-BC individuals (cohort 2), from which 37 were cancer-free (CF) individuals and 13 were patients with other cancers (OC) of the genitourinary/gynecological type. The clinical information of cohort 2 is summarized in Supplementary Tables S7. Overall, 16 GSL glycan species were detected in the urine samples (Fig. 3A, Supplementary Tables S3, S8 and S9). The PCA analysis integrating over all GSL-intensities did not show distinct clustering between the BC patients’ group and non-BC individuals (Supplementary Fig. S8A). Notably, intensity levels of GSLs belonging to the neolactotetra-series, being sialyl nLc4, α6- sialyl nLc4, Lewis X pentasaccharide (Le^X^ penta), fucosyl nLc4 and A type 2 hexasaccharide (A type 2 hexa), positively correlate with nLc4 in BC (Supplementary Fig. S8B). Of these neolactotetra-series glycans, levels of nLc4, sialyl nLc4 and Le^X^ penta are significantly upregulated in urinary EVs derived from patients with BC compared to non-BC individuals (multiple Mann-Whitney test, Fig. 3A). In particular, levels of nLc4 are significantly increased between patients with BC and non-BC individuals by a factor of 4.3 (*P* = 0.0011). Significantly increased levels of nLc4 are also observed when comparing BC patients either with only CF individuals (4.0-fold, *P* = 0.0044) or with patients with other cancers (5.5-fold, *P* = 0.0037) (Fig. 3B). ROC curve analyses reveal that the AUC level of nLc4 for BC diagnosis are 0.749 in BC vs non-BC (95% CI: 0.592-0.907; sensitivity: 56.3%; specificity: 90.0%), 0.734 in BC vs CF (95% CI: 0.568-0.900; sensitivity: 50.0%; specificity: 97.3%) and 0.793 in BC vs OC (95% CI: 0.628-0.959; sensitivity: 75.0%; specificity: 76.9%) (Supplementary Table S10). Similar analyses for all other identified GSLs can be found in Supplementary Fig. S9.

**Figure 3.**
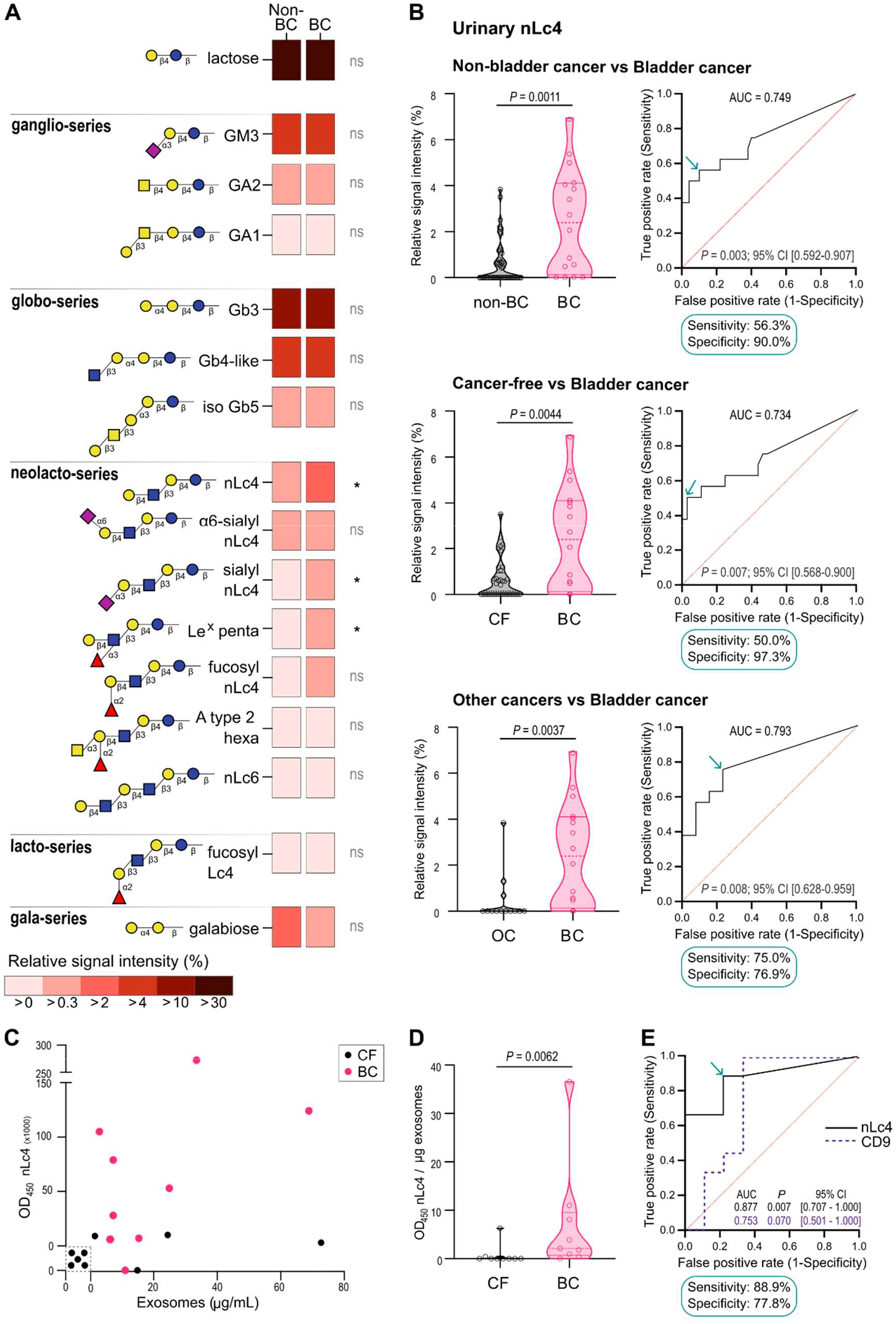
Glycosphingolipid profiling of urine liquid biopsies for bladder cancer biomarker discovery. GSLs detected in the urine of patients with bladder cancer (BC; n = 16) and without bladder cancer [non-bladder cancer (Non-BC; n = 50), comprising cancer-free individuals (CF, n = 37) and patients with other cancers (OC, n = 13)] through xCGE-LIF. **A,** Heatmap showing relative signal intensity mean values of 16 GSLs identified in BC. *Q* values were calculated using Multiple Mann-Whitney test (false discovery rate of 5%). *, *Q* value <0.05; ns, non-significant. **B,** Violin plots reporting the relative signal intensity of nLc4 among BC and non-BC (top), BC and CF (middle) and BC and OC (bottom). *P* values were calculated using two-tailed unpaired Mann-Whitney test; ROC curve analyses of urinary nLc4 for BC detection are shown (right). The arrows indicate the cut-off values of nLc4 relative signal intensity detected by xCGE-LIF that best discriminate bladder cancer detection. AUC, *P* values, 95% CI and optimal sensitivity and specificity values are shown. **C,** Correlation between nLc4 levels and CD9+ exosome concentration of urine samples from CF (n = 9) and BC patients (n = 9) as measured by ELISA. **D,** Violin plot showing the levels of nLc4 per μg of CD9+ exosomes derived from urine samples. *P* value was calculated using two-tailed unpaired Mann-Whitney test. **E,** ROC curve analyses for nLc4 and CD9 in patients with bladder cancer compared to healthy donors as predictive models for BC diagnosis. AUC, *P* values and 95% CI values are shown. The arrow indicates the cut-off value of nLc4 detected by ELISA that best discriminates bladder cancer detection (nLc4 cut-off value is expressed as OD_450_ x 1000). Optimal sensitivity and specificity values of nLc4 are shown. AUC, area under the curve; CI, confidence interval; Blue circle: glucose, yellow circle: galactose, blue square: *N*-acetylglucosamine, yellow square: *N*- acetylgalactosamine, pink diamond: *N*-acetylneuraminic acid, red triangle: fucose.

GSLs that are enriched in the urine of BC patients are likely to originate from the tumor or its microenvironment. For 13 BC patients of cohort 2 which were previously subjected to urinary xCGE-LIF analysis, we could additionally obtain FFPE tissue of the BC. We additionally performed xCGE-LIF analyses from these tissue samples and did a paired analysis of GSL- profiles from urine and tissue samples of the same BC patients (n = 13, Supplementary Fig. S10A, Supplementary Table S11). This analysis revealed that nLc4 levels are significantly increased in urine compared to the corresponding tumor tissues (Supplementary Fig. S10B), suggesting that nLc4 is preferentially secreted or shed by the malignant cells. To get to the bottom of this finding, we used the CAL-29 BC cell line to study secretion of GSLs *in vitro* and profiled GSL-glycans of cells and cell culture supernatant by xCGE-LIF. Similar to our findings on BC tissue samples of cohort 1, GM3, Gb3, Gb4-like and nLc4 are highly expressed by CAL- 29 cells (Supplementary Fig. S10C and D). Interestingly, from these GSLs, only nLc4 was detected in the supernatant fraction. These findings suggest that nLc4 is preferentially secreted/shedded by BC cells *in vitro* which is in line with our findings on BC patients’ urinary samples.

### Set-up of an ELISA enabling routine diagnostics detection of nLc4 as a bladder cancer urinary biomarker

To enable the clinical application of urinary EV-nLc4 as a diagnostic marker for bladder cancer without high end glyco-analytical instrumentation, we set-up a double sandwich ELISA for quantitative and qualitative analysis of urinary exosomes, without ultracentrifugation. To examine the correlation between nLc4 levels and CD9^+^-exosome concentration, EVs derived from 100 μL of urine were first captured with pan-exosome antibodies immobilized on a 96- well plate, in duplicate, and detected with either an anti-CD9 or 1B2 antibody. The ELISA system was used to analyze 18 urine samples from 9 patients with bladder cancer and 9 cancer-free individuals of cohort 2. The optical density of nLc4 measured for each urine sample was plotted against the corresponding exosome concentration (Fig. 3C). nLc4 expression is significantly higher in urinary EVs from patients with BC than in those from CF individuals (*P* = 0.0062; two-tailed unpaired Mann-Whitney test) (Fig. 3D). ROC curve analysis of nLc4 indicate that the AUC value for bladder cancer diagnosis is 0.877 (95% CI: 0.71-1.00), with a sensitivity of 88.9% and specificity of 77.8% (Fig. 3E).

## DISCUSSION

Bladder cancer is the most common malignancy of the urinary tract. Currently, cystoscopy with tissue biopsy and urinary cytology are the gold standard examination tools for BC diagnosis. However, because cystoscopy is an invasive procedure that can cause complications and cytology lacks adequate sensitivity and diagnostic efficacy (6, 19, 20), there is still an urgent need to develop new non-invasive and cost-effective tests for the reliable diagnosis of BC. Recent multi-omic studies have shown that the genomic, transcriptomic, proteomic and metabolomic profiles of bladder neoplasms allow for the stratification of BC patients into well-defined subtypes, providing as well a framework for biomarker discovery (21–29). The glycomic signature and, in particular, the glycolipidomic landscapes of bladder carcinomas, however, are still to be comprehensively explored. In fact, unraveling the GSL fingerprint of BC could provide considerable insights for future diagnosis and treatment targets, surpassing classical single GSL focused studies.

To date, most studies on GSL biomarker candidates for BC have been carried out in tissue samples and cell lines (30–32). Consequently, the extensive knowledge that can derive from liquid biopsies is yet to be unfolded. Here, we unravel for the first time the global GSL profile of both primary bladder tumor tissues, as well as urine samples from BC patients, showing that specific GSL structures are increased in BC.

Our study was enabled by the use of the xCGE-LIF workflow to globally decipher GSL glycosylation (15). This technique has particular advantages over other glyco-analytical technologies for analyzing patient samples, including the potential for high-throughput, low cost per sample, high sensitivity and even the capacity for isomer separation (33). Using this approach, we demonstrate that the overall GSL-signature of bladder cancer tissues is different from the one of cancer-free individuals. These results support the knowledge of specific GSLs being more highly expressed in tumors than in normal tissues (34). In particular, GM3, Gb3 and nLc4 glycans, the core precursors of the ganglio, globo-and neolacto-series respectively, are significantly increased in BC when compared to the correspondent paired normal adjacent tissues. This further corroborates the previously reported accumulation of GSL precursors in different human tumor models, an aberrant glyco-phenotype that results from the incomplete synthesis of certain GSLs (35–39). In BC specifically, GM3 and Gb3 are among the major GSLs that were identified so far in human bladder tumor tissue samples, being GM3 particularly increased in superficial tumors and Gb3 in invasive ones (30). In the present study, there are no differences regarding tumor classification as invasive or non-muscle invasive BC. Nevertheless, future studies comprising larger cohorts are needed to further correlate the expression of specific GSLs with the different pathological stages of bladder cancer.

Focusing on nLc4, the results are particularly interesting: besides being significantly increased in BC tissues, nLc4 is completely absent from both cancer-free and normal adjacent tissue sample groups. In fact, *B4GALT3* and *B4GALT4* (both genes encoding glycosyltransferases that generate nLc4 by adding a galactose to the terminal GlcNAc of the core Lc3 glycan in the β1-4 position) are upregulated in BC when compared to normal adjacent tissue samples of The Cancer Genome Atlas bladder cancer (TCGA-BLCA) dataset. This could explain why nLc4 is accumulated in bladder tumors while being completely absent from the normal tissue surroundings.

So far, the expression of nLc4 in human cancers has not been described in detail, although one study reports a higher level of nLc4 in the bone marrow of patients with acute myeloid leukemia when compared with healthy donors (40). Therefore, to our knowledge, this is the first time that nLc4 is identified as a cancer-specific GSL in a solid tumor. As xCGE-LIF does not provide information regarding the spatial distribution of the identified GSLs within the tumor microenvironment, we evaluated nLc4 as well as GATA3 expression in BC and control samples using specific monoclonal antibodies. The selective expression of GATA3 in a limited number of tissue types, mainly in the urothelium and breast glands, makes it a useful marker for BC, supporting the distinction of urothelial cancer from other genitourinary/gynecological diagnostic options (41, 42). As expected, the stained tissues show nuclear or cytoplasmic GATA3 positivity of bladder tumor cells (43), as well as a strong nuclear staining of the normal urothelial mucosa in the control tissue (44, 45). On the contrary, nLc4 expression is only observed on tumor cells, while all cell types within the normal bladder urothelium control are negative. Such observations become relevant when exploiting possible therapeutic applications, including chimeric antigen receptor (CAR) T-cell therapy, cancer vaccines, oncolytic viruses or monoclonal antibodies (46).

Focusing on the urine cohort (cohort 2), while the global GSL-signatures determined by xCGE-LIF are not distinct enough to discriminate between patients with BC from non-BC ones, the levels of nLc4 can correctly diagnose BC patients with an overall sensitivity of 56% and specificity of 90%. This finding is particularly relevant knowing that urine cytology, the most reliable test among urinary biomarker tools for detecting exfoliated tumor cells in urine, shows an overall sensitivity and specificity of up to 48% and 86%, respectively (47–49). Interestingly, the majority of the urinary GSL compounds identified in our study enclose the neolacto core chain, being almost all forms of this family positively correlated with their glycan precursor nLc4. Particularly, the Le^X^ pentasaccharide and nLc4 itself are significantly increased in samples derived from BC patients when compared to not only cancer-free individuals but also to patients with other genitourinary/gynecological malignancies. This is of relevance knowing that BTA *stat*^TM^, an FDA-approved test for BC, shows low specificity (46%) in subjects with other benign or malignant genitourinary diseases other than BC, giving rise to undesirable false positive results (50). Therefore, this finding could encompass an important advantage over other urinary biomarkers for the diagnosis of BC.

Previous studies have shown the immunostaining of Le^X^ antigen in exfoliated cells of urine samples to be a potentially useful diagnostic test for BC (51). However, until now there were no studies even reporting the presence of nLc4 in the urine of BC patients. In this direction, we evaluated next the relative abundance of specific GSLs, comparing both tissue and urine samples from the same BC patients. Notoriously, nLc4 presence is significantly increased in the urine of these patients, implying its possible shedding from the solid tumor. Moreover, the xCGE-LIF analysis of the cells and supernatant of the BC cell line used in this study shows that nLc4 is being preferentially secreted. These observations are in line with the previously described phenomenon of specific GSLs to be actively released from the membrane of tumor cells, being able to bind and influence the function of immune cells and, consequently, allowing tumor immune escape (52). Altogether, these results allow us to propose a new biosynthetic model for BC-specific differences in GSL glycosylation (Supplementary Fig. S11) and highlight the importance of further studies to unravel the mechanistic role of nLc4 shedding in BC. Finally, to evaluate the realistic prospects of urinary nLc4 clinical application as a biomarker for BC diagnosis, we did an ELISA for the quantitative and qualitative analysis of urinary exosomes, using both anti-CD9 and anti-nLc4 antibodies. With this test, the levels of exosomal nLc4 present in as few as two drops of urine (100 μL) could diagnose BC with overall sensitivity and specificity values of 89% and 78%, respectively. Being user friendly and cost-effective, this strategy displays advantages concerning other FDA-approved tests such as UroVysion^TM^, which is expensive, time-consuming and requires specialized laboratory equipment (47, 53, 54). Still, large-scale prospective cohort studies are invaluable to further test the robustness of nLc4 as a biomarker for BC in the future.

In conclusion, our work unveils for the first time the GSL-fingerprint of BC tissue and urine samples, highlighting the potential use of nLc4 as a therapeutic target and biomarker for diagnosis.

## METHODS

### Patient selection and sample collection

Paired primary tumor and adjacent normal bladder tissue from 30 patients at the Hannover Medical School were selected for cohort 1. Additionally, 7 normal bladder samples from cancer-free individuals were included. For cohort 2, urine samples were collected from 16 patients with BC and 50 non-BC individuals, from which there were 37 cancer-free individuals and 13 patients with other genitourinary/gynecological malignancies. Characteristics of samples and patients from both groups are listed in Supplementary Tables S1 and S2 (cohort 1) or S6 and S7 (cohort 2). Our study was conducted in accordance with the guidelines of the Hannover Medical School Ethics Committee (8619_BO_S_2019 and 10183_BO-K_2022). Samples were processed for xCGE-LIF analysis as described in the Supplementary Methods.

### Multiplexed capillary gel electrophoresis coupled to laser-induced fluorescence detection (xCGE-LIF)

Glycolipid extraction and xCGE-LIF analysis was performed as previously described (15). Briefly, deparaffinized tissue homogenates and urinary-EVs pellets were resuspended in 1 mL chloroform/methanol (1:2, v/v) and sonicated at 50% output in an ultrasonic bath for 15 min. Cell debris and proteins were removed by centrifugation and the supernatant was transferred into a glass vessel. The extraction was repeated twice using chloroform/methanol (2:1, v/v) and chloroform/methanol (1:1, v/v), sequentially. The pooled extracts were further purified on a Chromabond C_18_ ec polypropylene column (Macherey-Nagel, Dueren, Germany). The extracted glycolipids were digested with LudgerZyme Ceramide Glycanase (CGase) from *H. medicinalis* (Ludger, Abingdon, UK). Released glycans were fluorescently labeled with 8- aminopyrene-1,3,6-trisulfonic acid (APTS; Sigma-Aldrich, Germany) and excess of APTS and agents were removed by hydrophilic interaction liquid chromatography-solid phase extraction (HILIC-SPE). Finally, labeled glycans were separated and monitored by xCGE-LIF. The analysis was carried out using a remodeled ABI PRISM 3100-Avant Genetic Analyzer (ThermoFisher Scientific, Foster City, CA, USA). Data were assessed and processed using the GeneMapper Software (v.3.7.; Applied Biosystem, USA) and glycans were annotated using our in-house migration time database comprising 71 glycans. Quantification was performed based on relative fluorescence signal intensities, calculated for individual peak intensities (heights) in relation to the sum of all peak intensities.

### Multiplex immunofluorescence (mIF)

Staining of formalin-fixed paraffin embedded (FFPE) tissue was performed on deparaffinized sections. Heat-induced antigen retrieval was carried out in EDTA-based buffer (pH 9.0) at 98°C for 30 min, followed by washing steps in ice-cold water and in PBS. Sections were incubated with protein blocking solution (Zytomed Systems, Berlin, Germany) for 10 min at RT in a humid chamber, washed in PBS and incubated with primary antibodies specific to nLc4 (1:100, 1B2 hybridoma, (55) kindly provided by Ulla Mandel, Copenhagen Center for Glycomics) and GATA3 (1:300, monoclonal from rabbit, Abcam EPR16651, Cambridge, U.K) at 4°C, overnight. After a 3 min wash in PBS, samples were incubated with secondary antibodies, goat anti-mouse IgM Alexa Fluor® 488 (nLc4) and goat anti-rabbit IgG Alexa Fluor® 555 (Invitrogen, Thermo Fisher Scientific, Waltham, MA, U.S.A) 1:500 diluted in fluorescence antibody diluent (Zytomed Systems), for 30 min at RT in the dark. Sections were washed in PBS and mounted with Fluoromount-G^TM^ mounting medium with DAPI (Invitrogen, Thermo Fisher Scientific). Slides were examined and images acquired using a Zeiss Observer.Z1 microscope equipped with an AxioCam MRm digital camera. Afterwards, identical sections were stained with haematoxylin & eosin (H&E) and images of matched stained sections were acquired. Image analysis was performed with Image J software.

### Enzyme-linked immunosorbent assay (ELISA)

ELISA was performed to detect nLc4 on exosomes from urine samples using the ExoTEST^TM^ (Hansa BioMed, Estonia). Briefly, frozen urine samples from BC patients (n=9) and CF individuals (n=9) were thawed. Exosome standard solutions (serial dilution) and 100 μL of urine samples (in duplicate) were added to an ELISA plate pre-coated with pan-exosome antibodies for specific capture of exosomes. Samples were incubated overnight at 37°C and each well was washed by washing buffer three times, followed by incubation with either anti-CD9 (1:500; Hansa BioMed, Estonia) or anti-nLc4 (undiluted; (55)) antibodies for 2h at 37°C. After washing, samples were incubated with HRP-conjugated anti-mouse secondary antibodies (1:2000) for 1h at 37°C. The plate was developed with substrate chromogenic solution and stopped with stopping buffer (Hansa BioMed, Estonia). The absorbance was read at 450 nm with a microplate reader (Biotek PowerWave 340, BioTek Instruments, USA). The plate was also read at 570 nm and the measurement was subtracted from the measurement at absorbance 450 nm, following the manufacturer’s instructions. A calibration curve was interpolated with a linear regression model and the optical density of nLc4 measured for each urine sample was plotted against the correspondent exosome concentration.

### Statistical analyses

All statistical analyses were performed using the GraphPad Prism software (v9; GraphPad Software, CA, USA), the statistical software SPSS (v28.0.1.1; IBM Corp., IBM SPSS Statistics for Windows, NY, USA) or R programming language (v4.1.3; R Foundation for Statistical Computing, Vienna, Austria; http://www.r-prpject.org). Differences between groups were tested using the Multiple Mann-Whitney test (false discovery rate of 5%), the two-tailed unpaired Mann-Whitney test or the two-tailed Wilcoxon matched-pairs signed rank test. In ROC analyses, sensitivity and specificity were assessed as described in the Supplementary Methods.

## Authors’ Contributions

**I.B. Moreira**: Data curation, Formal Analysis, Investigation, Methology, Validation, Visualization, Writing – original draft. **C. Rossdam**: Investigation, Methology, Supervision. **J. Beimdiek**: Investigation, Methology. **M.M. Vicente**: Data curation, Investigation. **J. Schmitz:** Resources, Investigation. **A. Oberbeck**: Investigation. **U. Mandel**: Resources. **J.H. Bräsen**: Resources, Investigation. **H. Tezval**: Resources. **F.F.R. Buettner**: Conceptualization, Funding acquisition, Project administration, Supervision, Writing – original draft

## Acknowledgements

The authors would like to thank Prof. Dr. Rita Gerardy-Schahn, former head of the Institute of Clinical Biochemistry, Hannover Medical School (MHH) for providing an inspiring research atmosphere and general laboratory equipment. We also would like to thank E. Christians for expert technical assistance. We further thank Ulla Mandel, University of Copenhagen, Copenhagen Center for Glycomics for providing the hybridoma expressing the 1B2 antibody detecting nLc4. Thanks to all the subjects that accepted participating in this study and all the clinical staff involved in sample collection. We also acknowledge Sara Vicente for the support on the graphical design of schematic figures. This work was funded by the Deutsche Forschungsgemeinschaft (DFG, German Research Foundation) for Forschungsgruppe FOR2953 (Projektnummer: 409784463, project P9; BU 2920/4-2 for F.F.R.B.), for Forschungsgruppe FOR2509 (Projektnummer: 289991887, project P3; BU 2920/2-2 for F.F.R.B.) and for the Cluster of Excellence REBIRTH (From Regenerative Biology to Reconstructive Therapy, EXC 62), by the Lower Saxony Ministry of Science and Culture (Niedersächsisches Vorab) for the REBIRTH-Center, by the German Ministry for Education and Research (BMBF 13GW0399B for J.S. and J.H.B.), and by the Wilhelm Sander-Stiftung (for J.S. and J.H.B.)

## SUPPLEMENTARY METHODS

### Sample processing for xCGE-LIF

#### Tissue samples

Formalin-fixed paraffin embedded (FFPE) tissue samples were assessed by an expert pathologist that annotated tumor and healthy epithelial tissue regions from hematoxylin and eosin (H&E) stained slides by careful inspection under the microscope. From each tissue block, 5 mm biopsy punch samples were collected from both the tumor and normal tissue compartments and deparaffinized using xylene. Tissue samples were homogenized in chloroform/methanol (1:2, v/v) using the Precelly 24 tissue homogenizer with Precellys lysing kit (Bertin Instruments, Montigny-le-Bretonneux, France) 2 times for 10 seconds at 6500 rpm. Tissue homogenates were directly used for glycolipid extraction.

#### Urine samples

Small EVs were isolated from urine samples by differential centrifugation as previously described (56). Briefly, frozen urine samples (20 mL) were incubated at room temperature until they were partially thawed and vortexed. After complete thawing, samples were centrifuged at 200 g for 20 min at 4°C to remove cells, followed by centrifugation at 2000 g for 20 min at 4°C to eliminate cell debris. The supernatant was ultracentrifuged at 12,500 rpm for 20 min at 4°C using a 70Ti rotor (Beckman Coulter, CA, USA) to remove large EVS such as apoptotic bodies and microvesicles larger than 220 nm. The supernatant was filtered through a 0.22 μm filter. The filtrate was transferred to a new tube (Thickwall Polycarbonate Tube; Beckman Coulter, CA, USA) and ultracentrifuged at 34,200 rpm for 70 min at 4°C to isolate exosomes and small EVs. The pellet was then washed with PBS and ultracentrifuged at 26,500 rpm for 70 min at 4°C using a SW 32Ti rotor (Beckman Coulter, CA, USA). The final pellet was saved at -20°C until glycolipid extraction.

### Cell lines

#### Cell culture

The urinary bladder transitional cell carcinoma cell line CAL-29 (grade IV, stage T2) (57) was obtained from the DSMZ collection. The CAL-29 UGCG KO models was established using the CRISPR/Cas9 system (see Supplementary Methods). Cells were cultured in Dulbecco’s modified Eagle’s medium: nutrient mixture F-12 (DMEM/F12) (1:1) (PAN-Biotech, Aidenbach, Germany) supplemented with 10% fetal calf serum (FCS). All cells were grown at 37°C in an atmosphere of 5% CO_2_.

#### Sample processing for xCGE-LIF

Cells were grown at 90% confluency and 3 x 10^6^ cells were harvested, pelleted through centrifugation, washed twice with PBS and saved at -20°C until glycolipid extraction.

### Validation pipeline for anti-nLc4 antibody specificity

#### Establishment of UGCG knockout (KO) CAL-29 cells

The generation of *UGCG* KO of the cell line CAL-29 was performed using the CRISPR/Cas9 system as reported before (58). Briefly, sgRNA strands targeting exon 2 of human *UGCG* were cloned in the gRNA_AAVS1- T2 plasmid (Addgene #41818). The following gRNA strands were used: Top strand 5՛- CACCGTTAGGATCTACCCCTTTCAG-3՛; Bottom strand 5՛- AAACCTGAAAGGGGTAGATCCTAA-3՛. CAL-29 cells were transfected with the sgRNA-containing plasmid and the pCas9_GFP plasmid (Addgene #44719) using Nucleofector, were sorted based on GFP expression using FACSAria™ Fusion FACS sorter and further cultured in single cell clones. KO clones were screened by anti-GM3 ((clone GMR6; Tokyo Chemical Industry, Tokio, Japan) flow cytometry and immunofluorescence analysis, and mutations were confirmed by sequencing.

#### Immunofluorescence

Cells were seeded in 12-well plates coverslips and fixed with paraformaldehyde. Nonspecific binding was blocked with 5% BSA in PBS for 30 min at room temperature. Cells were stained with monoclonal antibody 1B2 (anti-nLc4; hybridoma kindly provided by Ulla Mandel, Copenhagen Center for Glycomics, (55)) for 1h at room temperature. Anti-mouse IgM Alexa Fluor® 488 secondary antibody was used at 1:500 for 1h at room temperature. Cells were washed with PBS and nuclear stain was performed with DAPI (VECTASHIELD® antifade mouting medium with DAPI, Vector laboratories, CA, USA). Images were acquired using a Zeiss Observer.Z1 microscope equipped with an AxioCam MRm digital camera. Image analysis was performed using the Image J software.

#### Flow cytometry

Cell surface nLc4 expression was analyzed using flow cytometry. The cells were incubated with undiluted anti-nLc4 antibody (55) for 30 min on ice, washed with 1% FCS in PBS and incubated with anti-mouse IgM Alexa Fluor® 488 secondary antibody (1:500) for 30 min on ice. Propidium iodide (1:1000, Sigma-Aldrich, Germany) was used as a viability dye. Cells were analyzed using in a CyFlow®xxx flow cytometer (Partec, Munster, Germany) and the data were analyzed with FlowJo v10 software.

### Gene expression datasetTCGA-BLCA data analysis

The transcriptomic data of the TCGA-BLCA cohort was downloaded using the TCGAbiolinks Bioconductor package (v2.25.3). Prior to analysis, preprocessing steps were done, data was normalized and differential gene expression between BC and NAT samples was done using the package DEseq2 (v1.34.0).

### Statistical analyses

The prediction capacity of all identified GSL markers for BC diagnosis was determined by plotting the receiver operating characteristic (ROC) curves and calculating the area under the curve (AUC). The relative signal intensity cut-off value that revealed the best balance between sensitivity and specificity, defined at the highest Youden’s index [*sensitivity* + *specificity* − 1], was selected for the subsequent statistical analysis. Sensitivity, specificity, 95% confidence intervals (CI) and *P* values were calculated. The prediction probabilities for the combination of GM3, Gb3 and nLc4 levels were calculated by performing a multivariable logistic regression model including these three variables, and a ROC curve analysis to assess the global predictive ability of the model and the sensitivity and specificity for different cut-off values. A principal component analysis (PCA) based on the relative signal intensity values of the identified GSLs was carried out to examine clustering patterns of BC samples and controls. Samples were plotted in a 2-D space consisting of two principal components (PC1 and PC2). The correlations between the levels of the GSLs identified were described for BC samples using Spearman rank correlation coefficient.

---

**Supplementary Figure S1.**
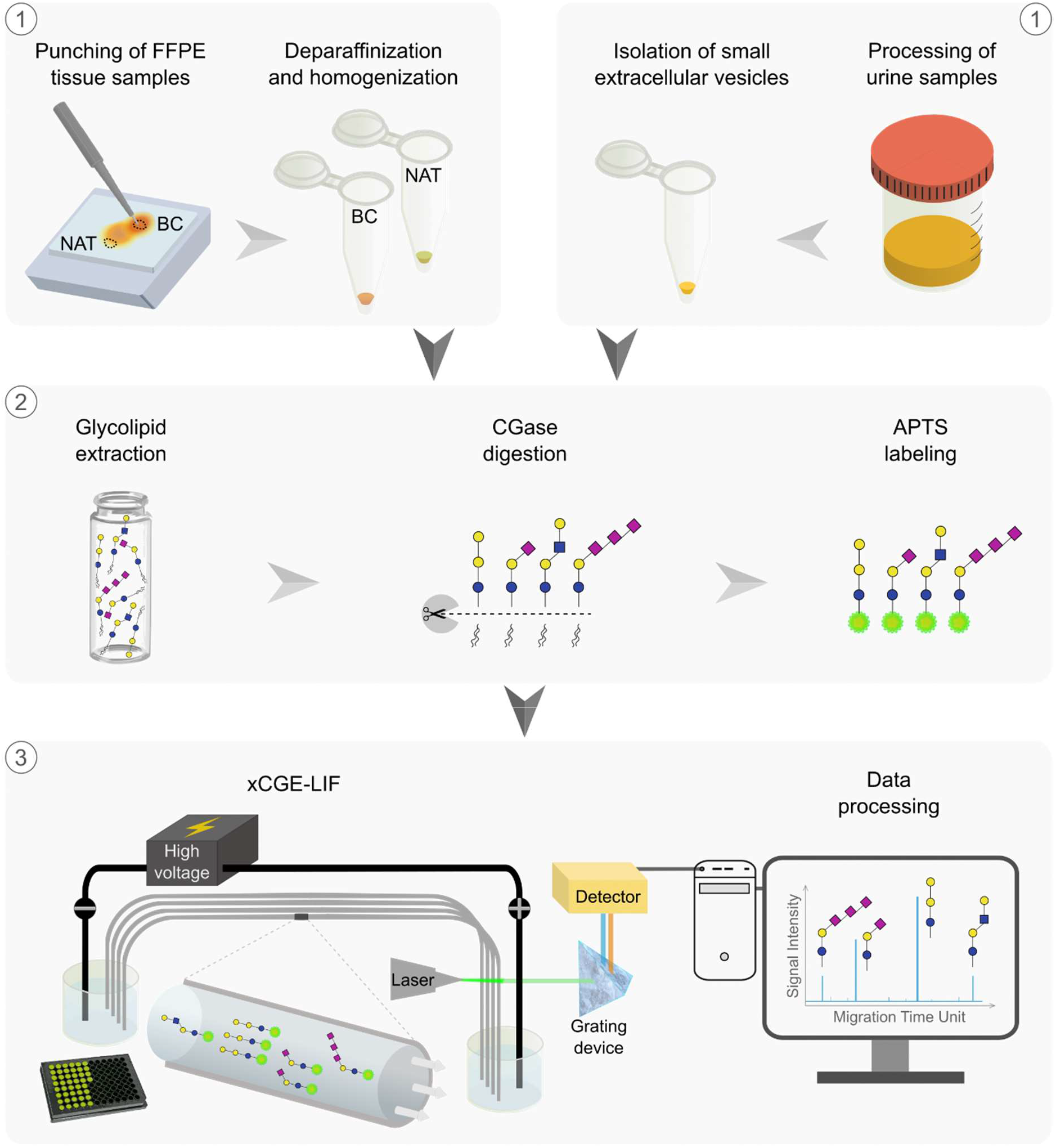
Schematic workflow for GSL-glycomics from punched FFPE tissues and urine samples. Bladder tumor and normal adjacent tissue (NAT) regions of FFPE blocks were selected, punched and deparaffinized. 20 mL of urine samples were processed for small extracellular vesicles isolation through sequential centrifugation and ultracentrifugation steps. The tissue lysates and urinary small extracellular vesicle pellets obtained were used for GSL extraction, CGase digestion, APTS fluorescent labeling and purification for xCGE-LIF analysis. Labeled glycans were separated and monitored by xCGE- LIF, data were processed and glycans were annotated.

**Supplementary Figure S2.**
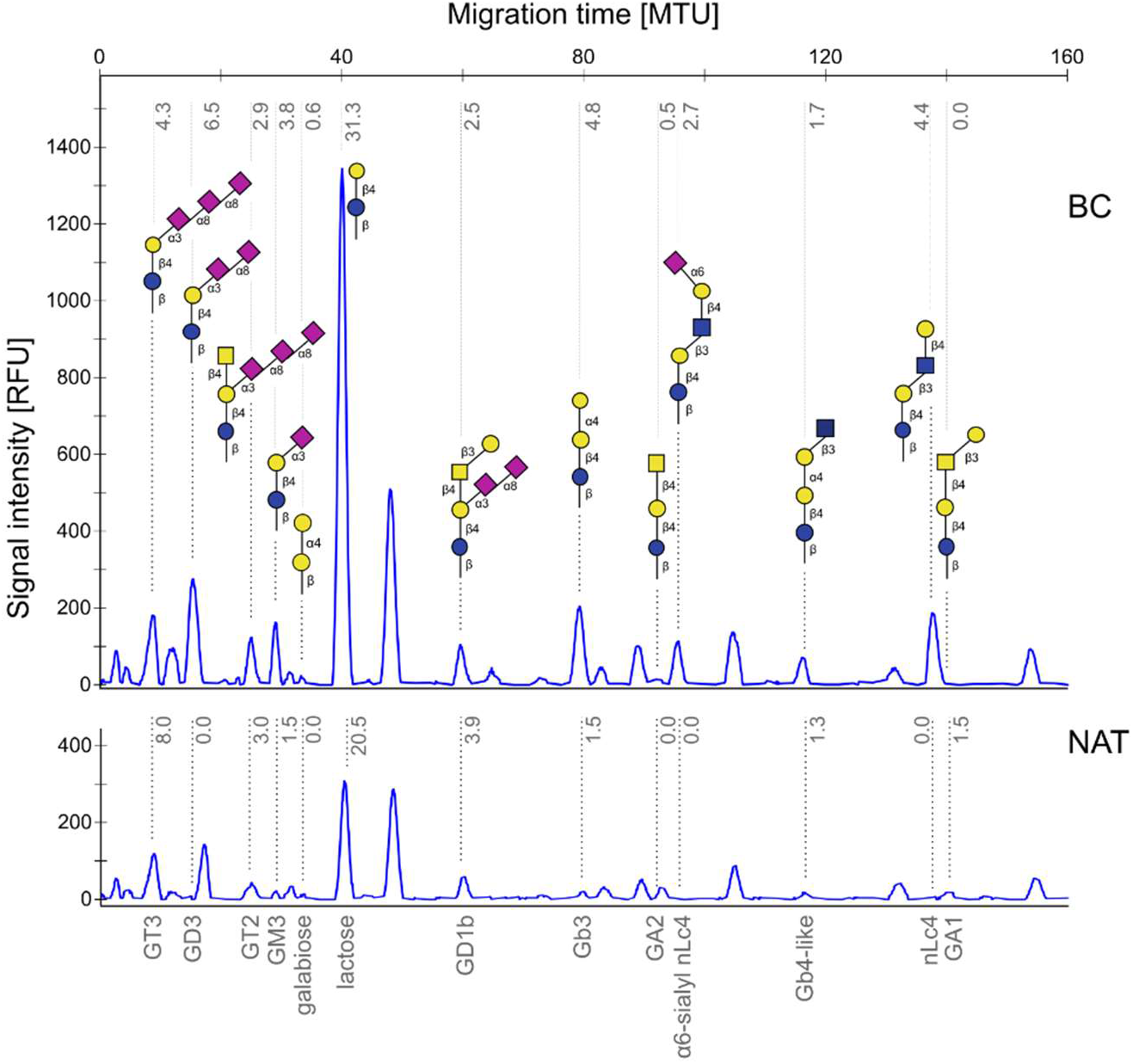
Example of GSL electropherogram profiles obtained from the bladder cancer and the normal adjacent tissue samples of patient number 24 (BC24, up; NAT24, down). Values in gray correspond to calculated relative signal intensities (in percentage). RFU, relative fluorescence units; MTU, migration time unit; Blue circle: glucose, yellow circle: galactose, blue square: *N*-acetylglucosamine, yellow square: *N*- acetylgalactosamine, pink diamond: *N*-acetylneuraminic acid.

**Supplementary Figure S3.**
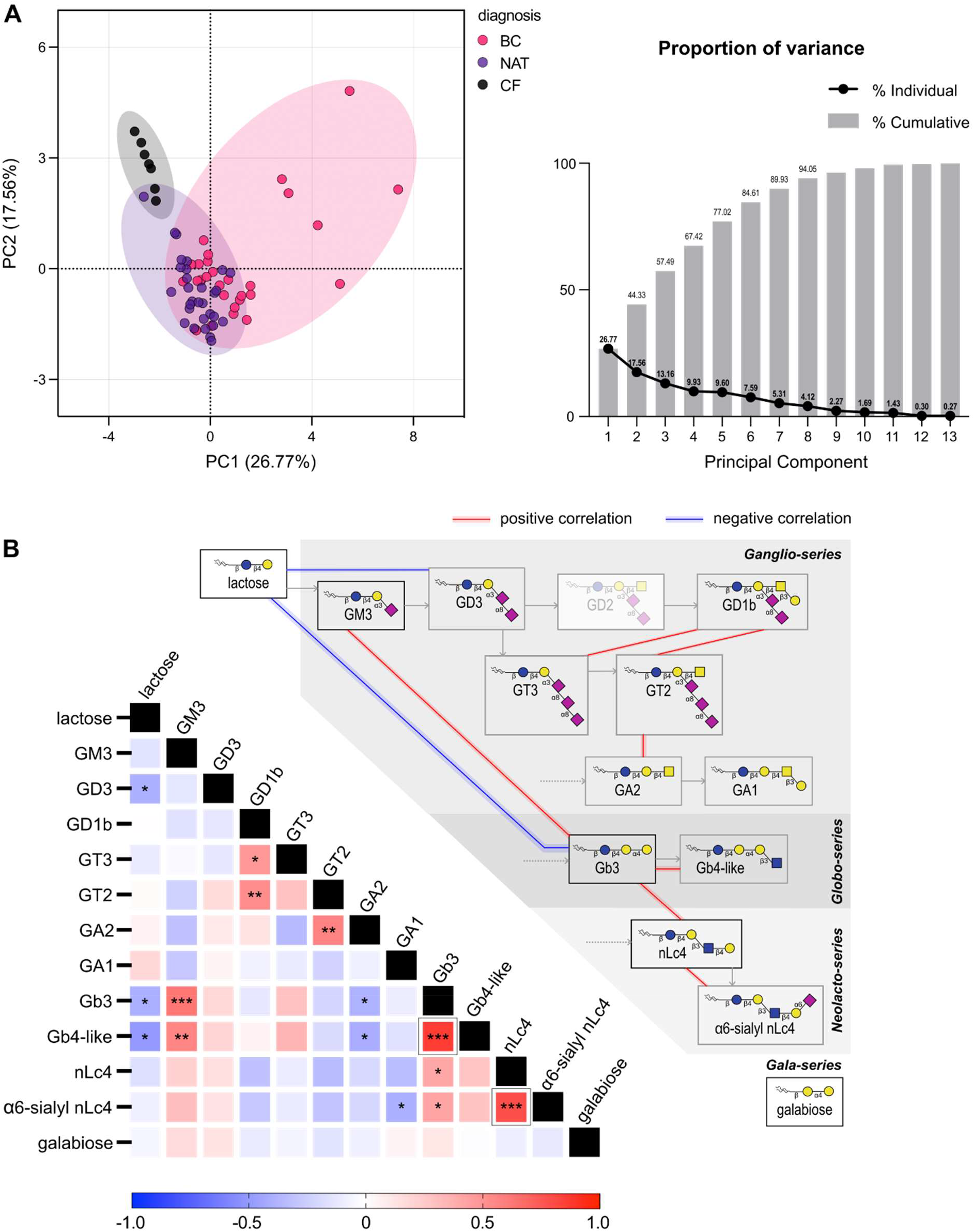
Analysis of tumor associated glycosphingolipids identified in bladder cancer tissue samples. **A,** Principal component analysis (PCA) model based on the relative abundance (%) of individual GSLs expressed in bladder tissues. Separation between cancer, normal adjacent tissue and cancer-free is illustrated on the left; Proportion of variance of the principal components is shown on the right. The top two principal components (PC1 and PC2) explain 44.3 % of the variation within the data. **B,** Correlation matrix of GSL signatures using Spearman correlation coefficients for bladder cancer samples (left). Black boxes highlight the positive correlation between Gb4-like/Gb3 and α6-sialyl nLc4/nLc4; Representation of the GSL biosynthesis pathway and the main correlation relationships are shown on the right. *, *P* value <0.05; **, *P* value <0.005; ***, *P* value <0.001; Blue circle: glucose, yellow circle: galactose, blue square: *N*-acetylglucosamine, yellow square: *N*- acetylgalactosamine, pink diamond: *N*-acetylneuraminic acid.

**Supplementary Figure S4.**
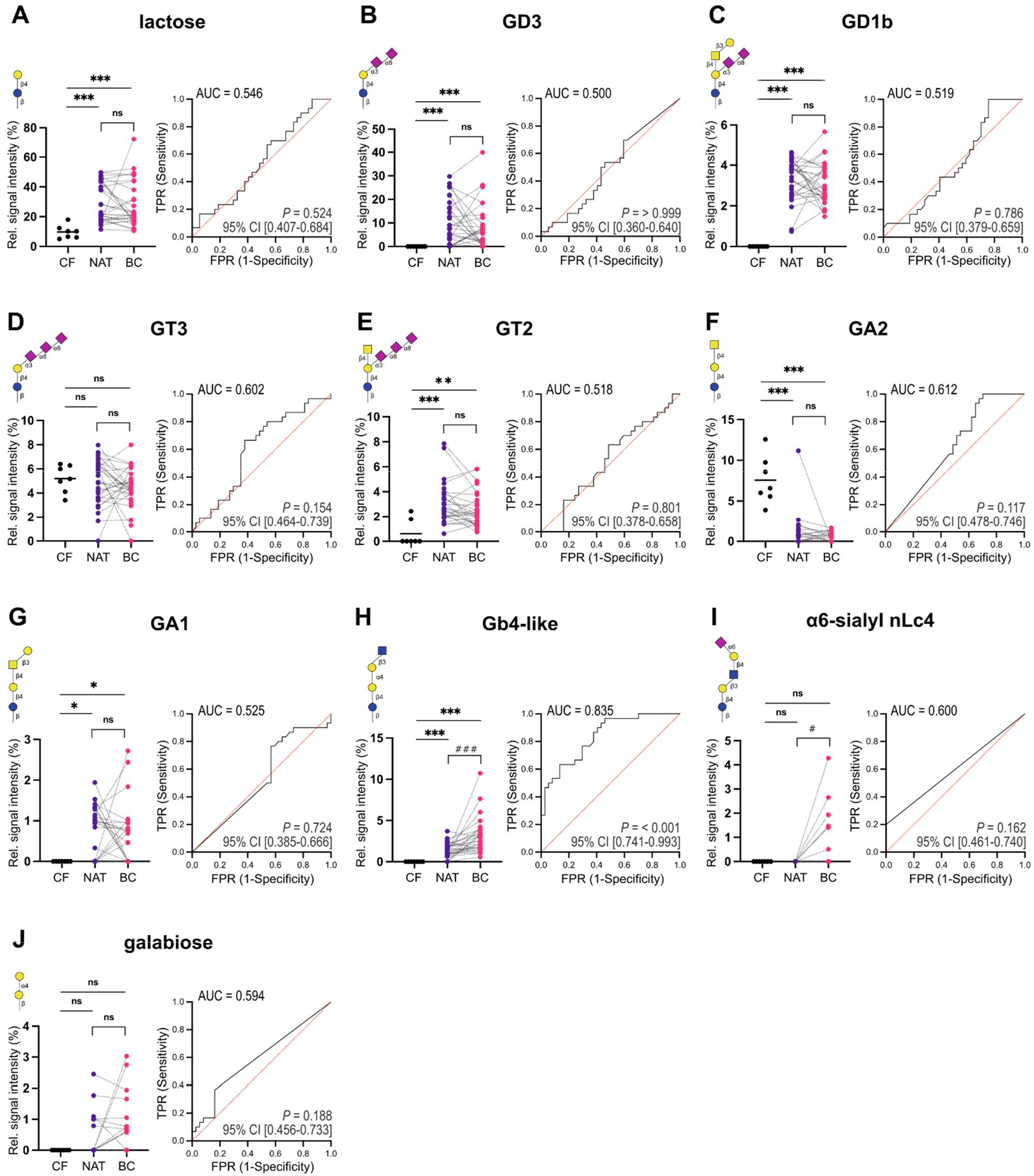
Glycosphingolipids detected in bladder cancer (BC, n = 30), normal adjacent tissue (NAT, n = 30) and cancer-free (CF, n = 7) tissue samples through xCGE-LIF. Relative signal intensity levels of **A,** lactose, **B,** GD3, **C,** GD1b, **D,** GT3, **E,** GT2, **F,** GA2, **G,** GA1, **H,** Gb4-like, **I,** α6-sialyl nLc4 and **J,** galabiose in CF and paired analysis between NAT and BC (left). *P* values were calculated using two-tailed unpaired Mann-Whitney test (for comparisons with the CF group) or two-tailed Wilcoxon matched-pairs signed rank test (for NAT *vs.* BC comparison); ROC curve analysis in bladder cancer detection obtained by calculating the sensitivity and specificity of the test at every possible cut-off point and plotting the sensitivity against 1-specificity are shown on the right. AUC, *P* value and 95% CI values are shown. *, *P* value <0.05; **, *P* value <0.005; ***, *P* value <0.001; ns, non-significant; AUC, area under the curve; CI, confidence interval; TPR, true positive rate; FPR, False positive rate; blue circle: glucose, yellow circle: galactose, blue square: *N*-acetylglucosamine, yellow square: *N*-acetylgalactosamine, pink diamond: *N*-acetylneuraminic acid.

**Supplementary Figure S5.**
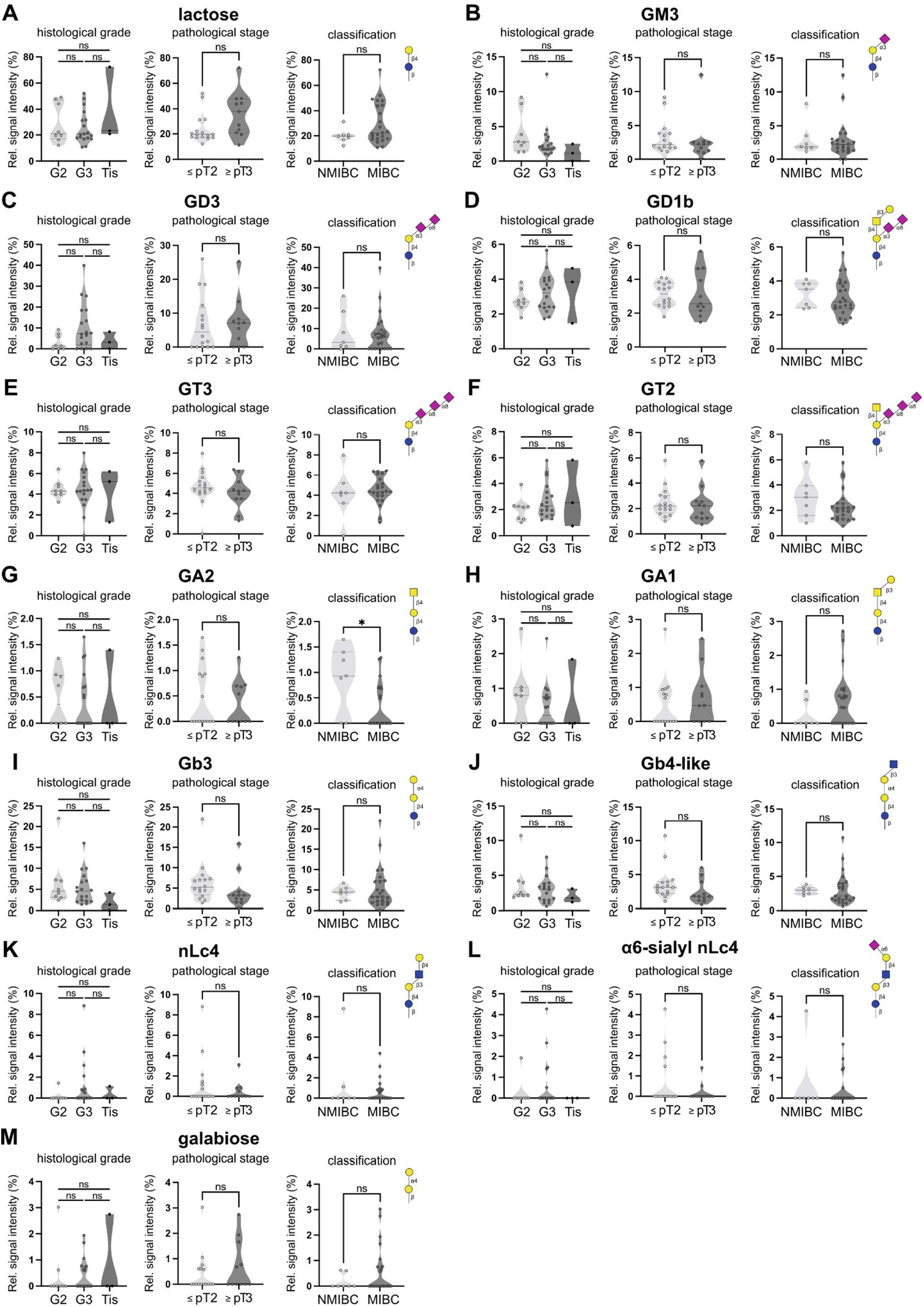
Glycosphingolipids detected in bladder cancer (BC, n = 30) tissue samples through xCGE-LIF. Relative signal intensity levels of **A,** lactose, **B,** GM3, **C,** GD3, **D,** GD1b, **E,** GT3, **F,** GT2, **G,** GA2, **H,** GA1, **I,** Gb3, **J,** Gb4-like, **K,** nLc4, **L,** α6-sialyl nLc4 and **M,** galabiose across disease grades (left), stage^a^ (center) and tumor classification (right). *P* values were calculated using two-tailed unpaired Mann-Whitney tests. G2, grade 2 (n = 8); G3, grade 3 (n = 18); Tis, tumor *in situ* (n = 3); Stage ≤ pT2 (includes pTa, pT1 and pT2; n = 16), stage ≥ pT3 (includes pT3 and pT4; n = 11); NMIBC, non-muscle invasive bladder cancer (n = 7); MIBC, muscle invasive bladder cancer (n = 23); *, *P* value <0.05; ns, non-significant; blue circle: glucose, yellow circle: galactose, blue square: *N*-acetylglucosamine, yellow square: *N*- acetylgalactosamine, pink diamond: *N*-acetylneuraminic acid. ^a^ individuals with cancer stage determined after neoadjuvant chemotherapy were excluded from the analysis.

**Supplementary Figure S6.**
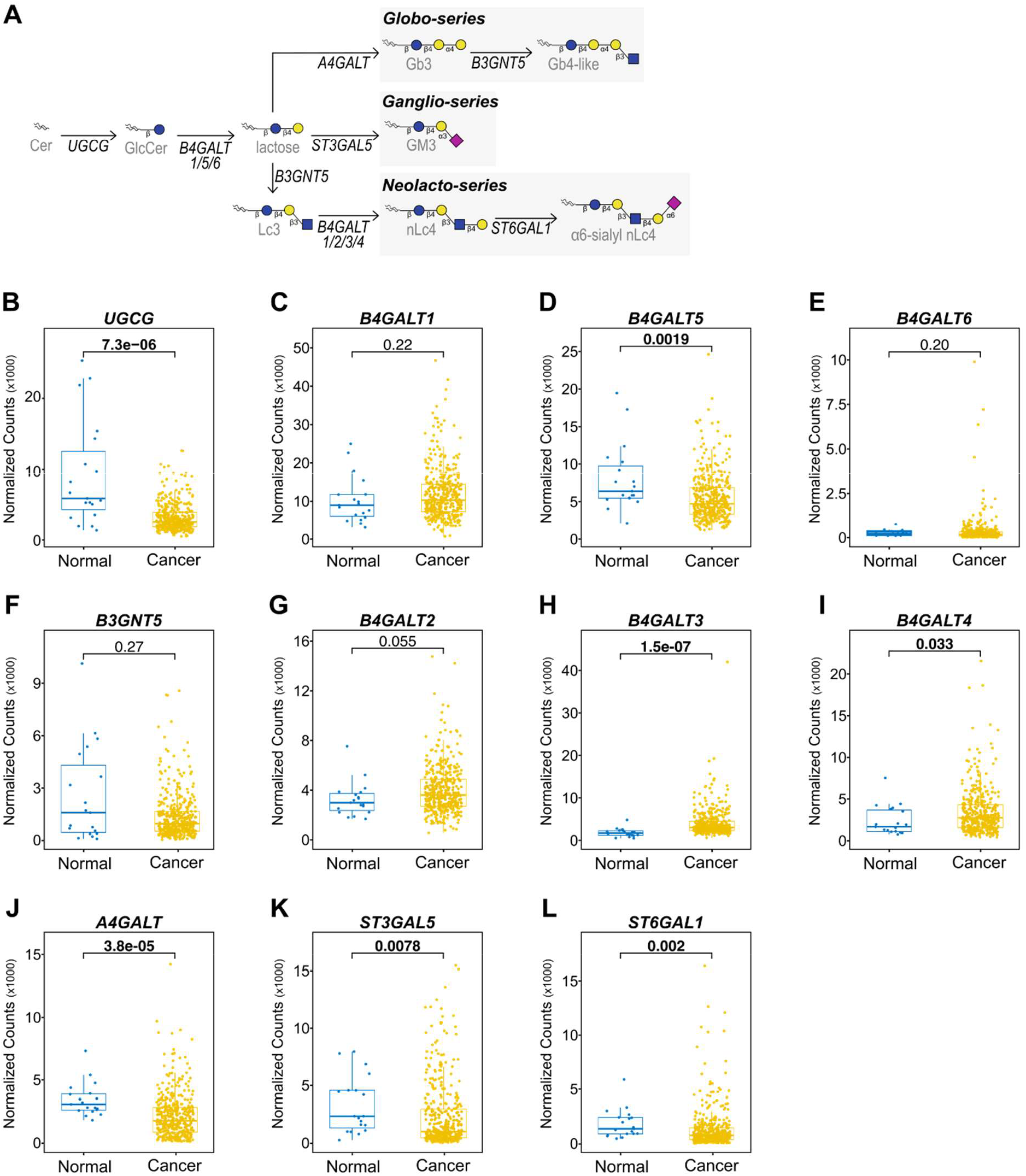
Comparison of gene expression of glycosyltransferases in bladder cancer and normal adjacent bladder. **A,** Biosynthetic pathways of major GSL-series. Differences in gene expression in cancer compared to normal surrounding bladder tissue are observed for **B,** UGCG, **C,** B4GALT1, **D,** B4GALT5, **E,** B4GALT6, **F,** B3GNT5, **G,** B4GALT2, **H,** B4GALT3, **I,** B4GALT4, **J,** A4GALT, **K,** ST3GAL5 and **L,** ST6GAL1 glycosyltranseferases. Raw data was obtained from the TCGA-BLCA (The Cancer Genome Atlas Urothelial Bladder Carcinoma) dataset. Blue circle: glucose, yellow circle: galactose, blue square: *N*- acetylglucosamine, yellow square: *N*-acetylgalactosamine, pink diamond: *N*-acetylneuraminic acid.

**Supplementary Figure S7.**
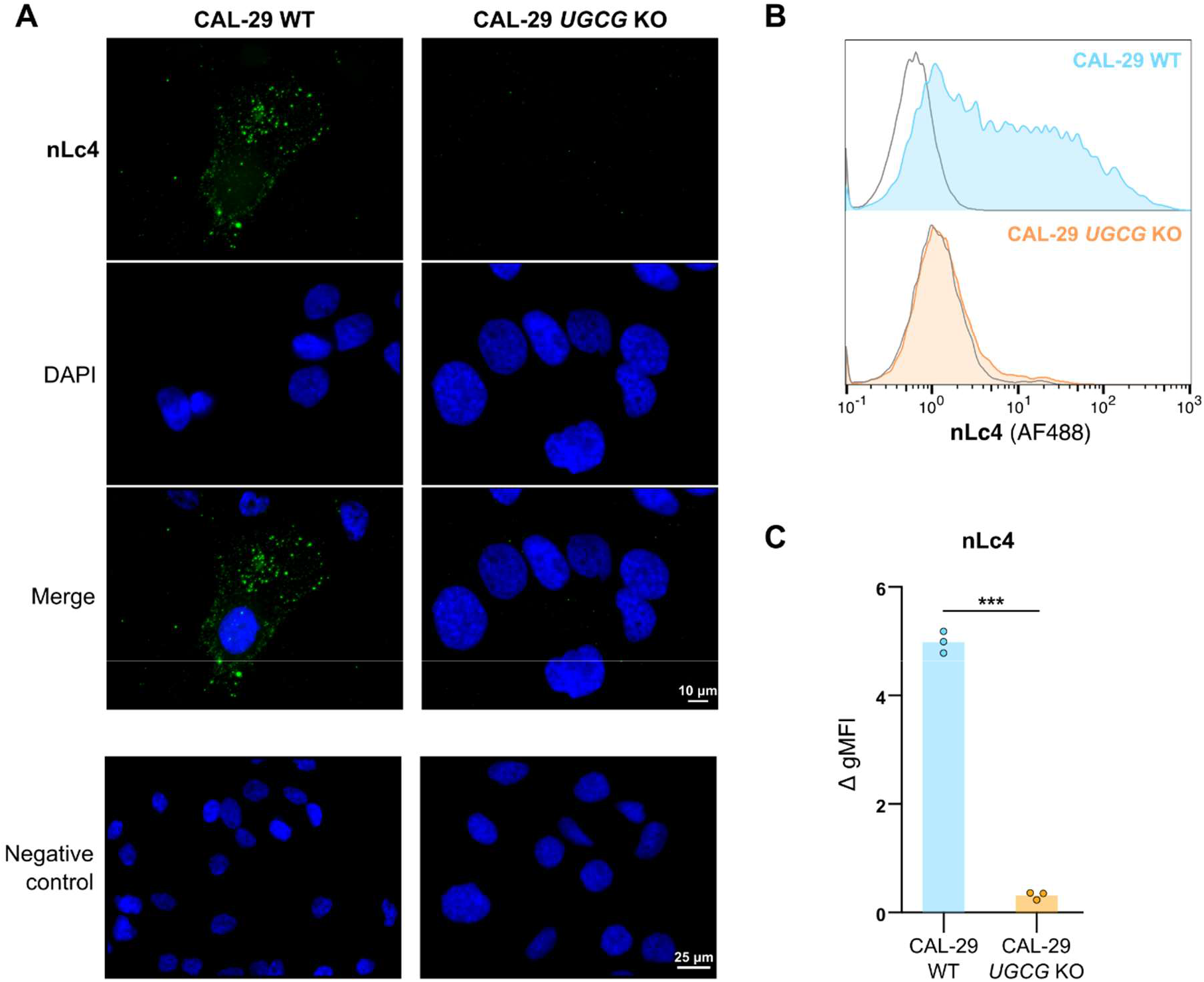
Validation pipeline for nLc4-specific antibody. **A,** Immunofluorescence analysis of nLc4 in a bladder cancer cell line model of glucose-derived GSL deficiency (CAL-29 *UGCG* KO) and its control counterpart (CAL-29 WT). Scale bar corresponds to 10 μM. Negative control scale bar corresponds to 25 μM. **B,** Histograms of nLc4 fluorescence signal in flow cytometry. Negative controls are shown in gray lines. **C,** Geometric mean fluorescence intensity (gMFI) quantification of fluorescence signal related to nLc4 expression in CAL-29 *UGCG* KO and CAL-29 WT cells (N = 3). All gMFI levels were subtracted the negative control intensity. *P* value was calculated using unpaired t test with Welch’s correction (Welch’s t test). ***, *P* value <0.001.

**Supplementary Figure S8.**
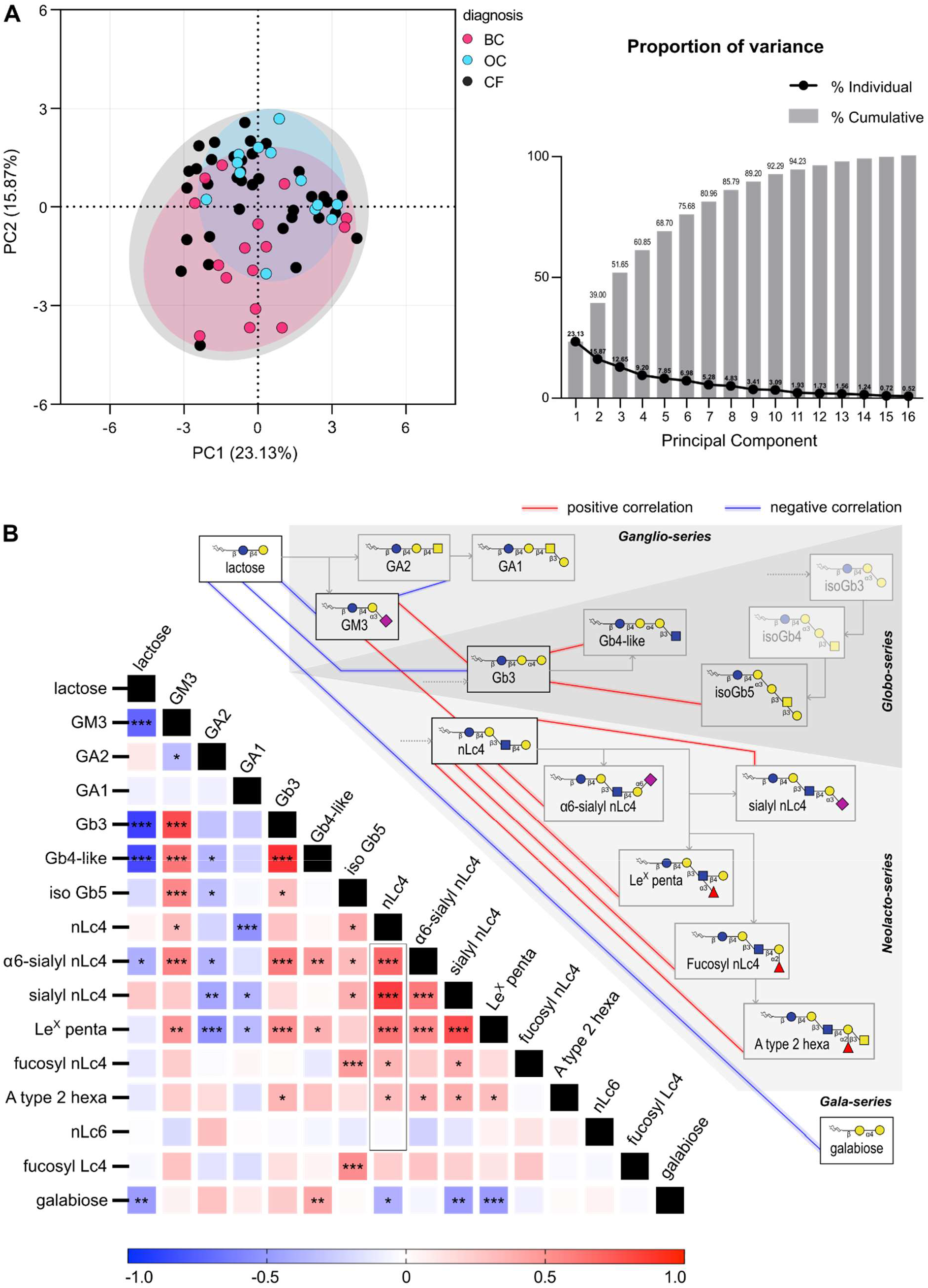
Analysis of tumor associated glycosphingolipids identified in bladder cancer urine samples. **A,** Principal component analysis (PCA) model based on the relative abundance (%) of individual GSLs expressed in urine samples. Separation between bladder cancer, other cancers and cancer-free is illustrated on the left; Proportion of variance of the principal components is shown on the right. The top two principal components (PC1 and PC2) explain 38.7 % of the variation within the data. **B,** Correlation matrix of GSL signatures using Spearman correlation coefficients for bladder cancer samples (left). Black box highlights the correlation between nLc4 and other GSLs of the Neolacto-series; Representation of the GSL biosynthesis pathway and the main correlation relationships are shown on the right. *, *P* value <0.05; **, *P* value <0.005; ***, *P* value <0.001; Blue circle: glucose, yellow circle: galactose, blue square: *N*-acetylglucosamine, yellow square: *N*-acetylgalactosamine, pink diamond: *N*-acetylneuraminic acid, red triangle: fucose.

**Supplementary Figure S9.**
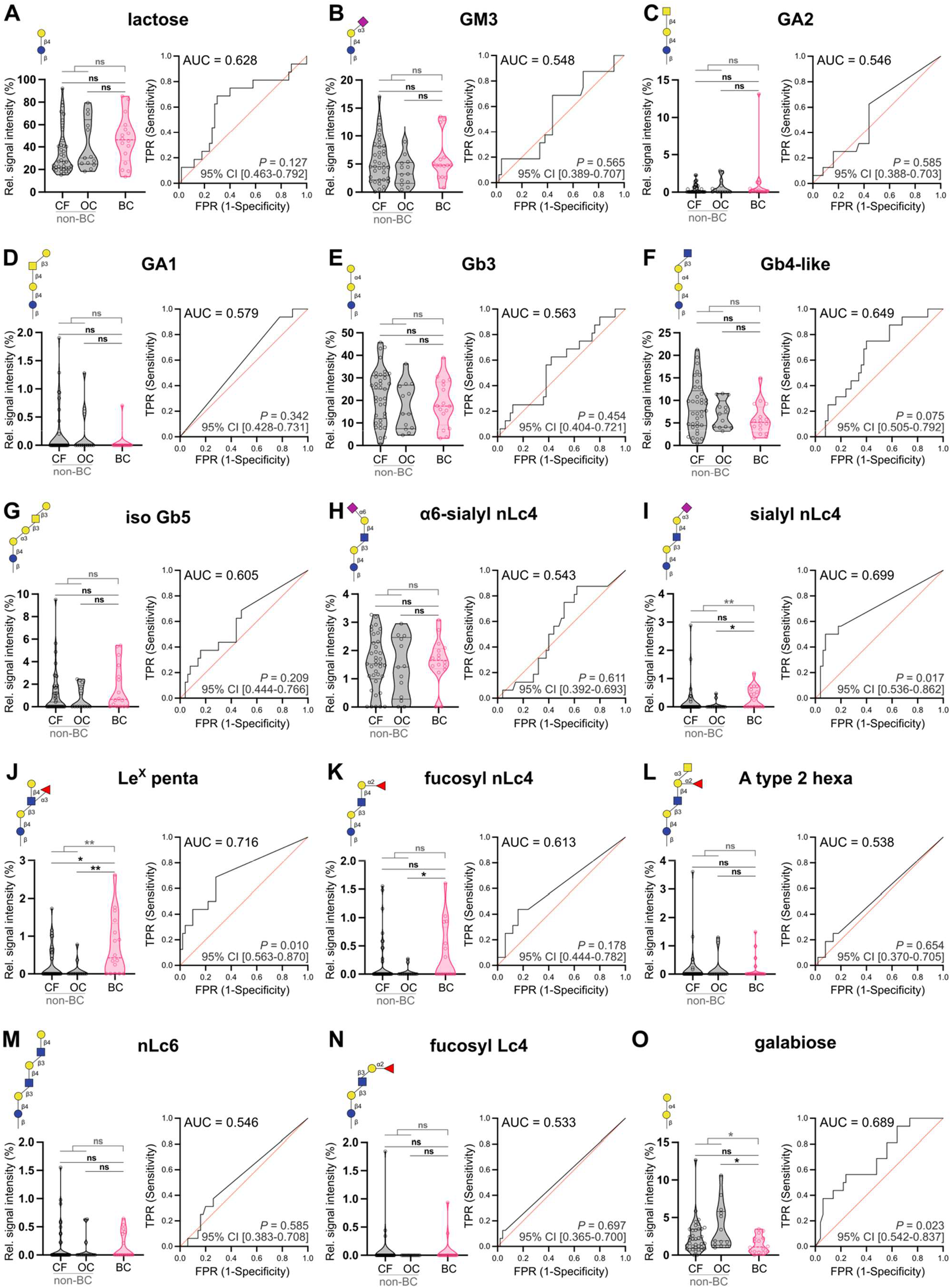
Glycosphingolipids detected in urine of patients with bladder cancer (BC; n = 16) and without bladder cancer [non-bladder cancer (Non-BC; n = 50), comprising cancer-free individuals (CF, n = 37) and patients with other cancers (OC, n = 13)] through xCGE-LIF. Violin plots reporting the relative signal intensity of **A,** lactose, **B,** GM3, **C,** GA2, **D,** GA1, **E,** Gb3, **F,** Gb4-like, **G,** iso Gb5, **H,** α6-sialyl nLc4, **I,** sialyl nLc4, **J,** Le^X^ penta, **K,** fucosyl nLc4, **L,** A type 2 hexa, **M,** nLc6, **N,** fucosyl Lc4 and **O,** galabiose (left). *P* values were calculated using two-tailed unpaired Mann-Whitney test; ROC curve analyses of urinary GSLs is patients with BC compared to non-BC individuals as predictive models for BC diagnosis (right). AUC, *P* values and 95% CI values are shown. *, *P* value <0.05; **, *P* value <0.005; ns, non-significant; AUC, area under the curve; CI, confidence interval; TPR, true positive rate; FPR, False positive rate; blue circle: glucose, yellow circle: galactose, blue square: *N*-acetylglucosamine, yellow square: *N*-acetylgalactosamine, pink diamond: *N*- acetylneuraminic acid, red triangle: fucose.

**Supplementary Figure S10.**
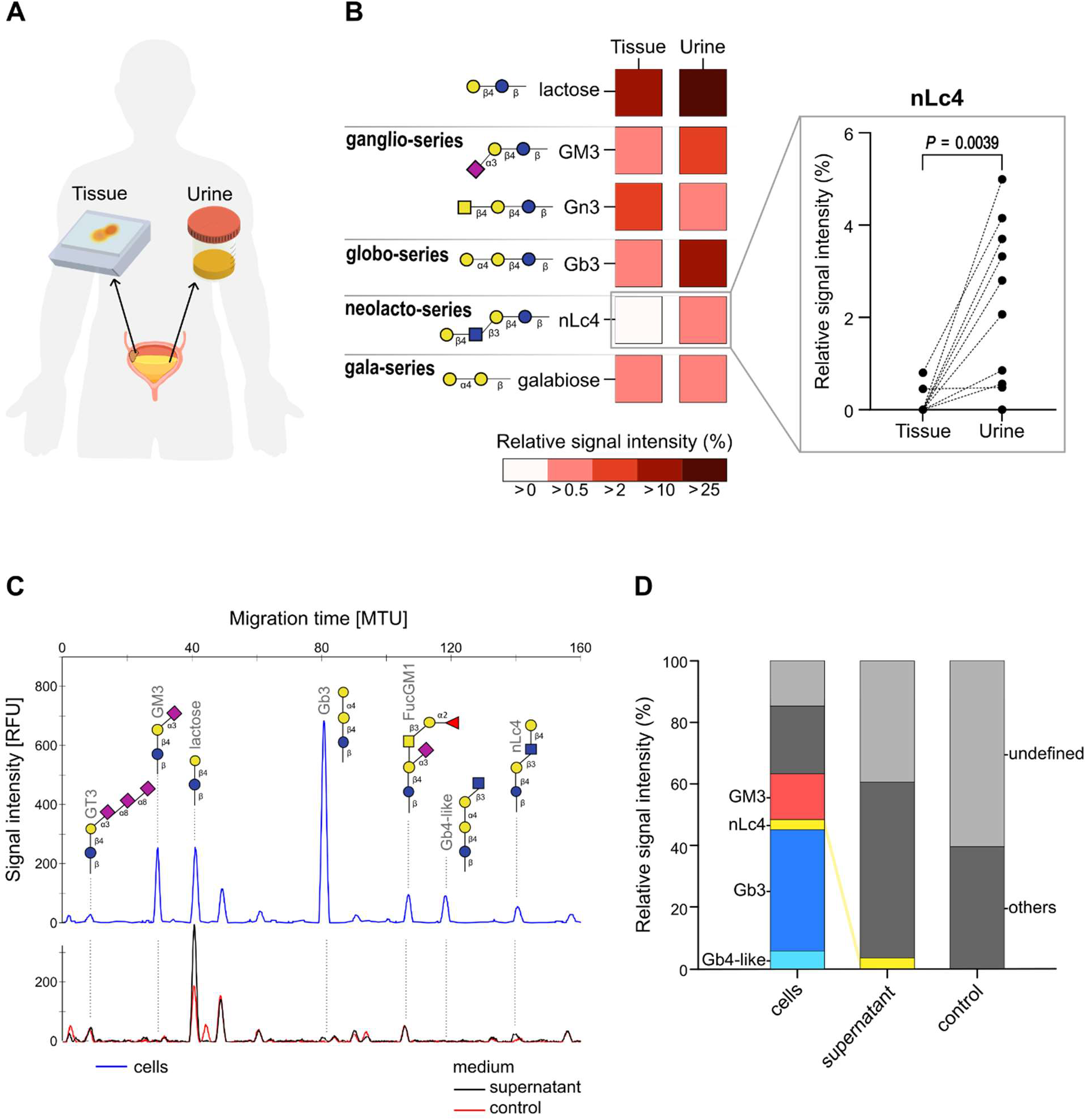
Glycosphingolipid secretion in bladder cancer. **A,** Schematic representation of tissue and urine paired sample collection from a BC patient (N = 13). **B,** Heatmap showing relative signal intensity mean values of 6 GSLs identified in both tissue and urine samples from patients with BC (left); Paired analysis of nLc4 signal between tissue and urine samples (right). *P* value was calculated using two-tailed Wilcoxon matched-pairs signed rank test. **C,** GSL electropherogram profiles obtained from CAL-29 WT cells (top, blue), CAL-29 WT supernatant culture medium (bottom, black) and culture medium only as control (bottom, red). **D,** Comparison between relative signal intensity values of the glycosphingolipids detected in CAL-29 WT cells, CAL-29 WT supernatant and cell culture medium only. RFU, relative fluorescence units; MTU, migration time unit; Blue circle: glucose, yellow circle: galactose, blue square: *N*-acetylglucosamine, yellow square: *N*-acetylgalactosamine, pink diamond: *N*-acetylneuraminic acid, red triangle: fucose.

**Supplementary Figure S11.**
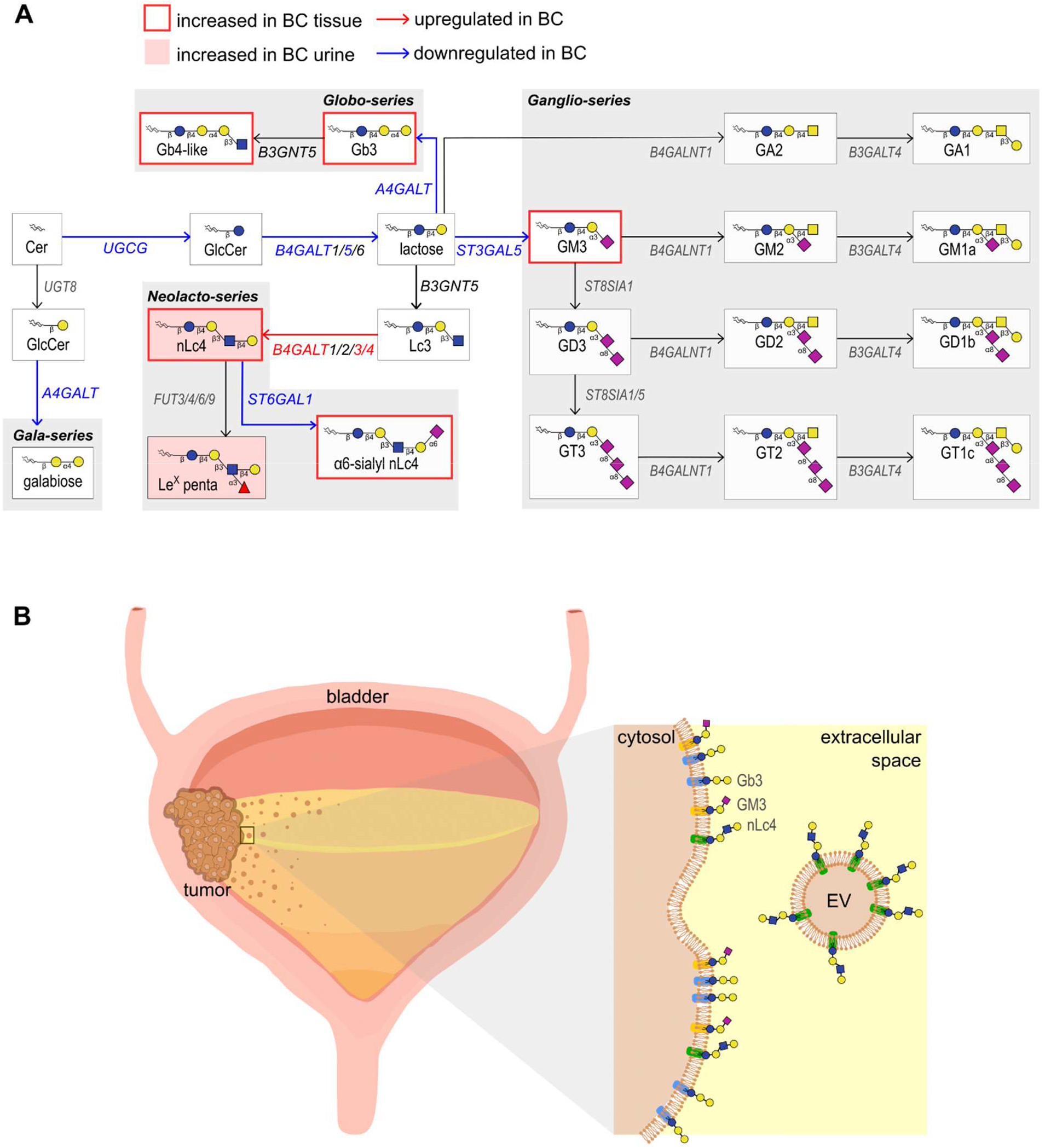
Bladder cancer specific glycosphingolipid alterations. **A,** Proposed biosynthetic model explaining the GSL changes that occur in bladder cancer by integrating GSL-glycomics (tissue and urine samples) with gene expression results. **B,** Proposed schematic representation of the preferential shedding of nLc4-containing-EVs by bladder tumor cells into the voided urine. EV, extracellular vesicle; Blue circle: glucose, yellow circle: galactose, blue square: *N*-acetylglucosamine, yellow square: *N*-acetylgalactosamine, pink diamond: *N*-acetylneuraminic acid, red triangle: fucose.

